# Molecular mechanism of N-terminal acetylation by the ternary NatC complex

**DOI:** 10.1101/2021.02.01.429250

**Authors:** Sunbin Deng, Leah Gottlieb, Buyan Pan, Julianna Supplee, Xuepeng Wei, E James Petersson, Ronen Marmorstein

**Author notes:** Correspondence (R.M.).

## Abstract

Protein N-terminal acetylation is predominantly a ribosome-associated modification, with NatA-E serving as the major enzymes. NatC is the most unusual of these enzymes, containing one Naa30 catalytic subunit and two auxiliary subunits, Naa35 and Naa38; and substrate specificity profile that overlaps with NatE. Here, we report the Cryo-EM structure of *S. pombe* NatC with a NatE/C-type bisubstrate analogue and inositol hexaphosphate (IP_6_), and associated biochemistry studies. We find that the presence of three subunits is a prerequisite for normal NatC acetylation activity in yeast and that IP_6_ binds tightly to NatC to stabilize the complex. We also describe the molecular basis for IP_6_-mediated NatC complex stabilization and the overlapping yet distinct substrate profiles of NatC and NatE.

## Introduction

Protein N-terminal acetylation is one of the most common protein modifications, occurring on 50-85% of eukaryotic proteins and 10–30% of bacterial and archaea proteins (Aksnes et al., 2016; Arnesen et al., 2009; Deng and Marmorstein, 2021). N-terminal acetylation changes the local chemical properties of protein N-termini and impacts diverse protein functions including stability, half-life, folding, aggregation, degradation, complex formation, and localization (Aksnes et al., 2016; Arnesen, 2011; Deng and Marmorstein, 2021; Dikiy and Eliezer, 2014; Forte et al., 2011; Holmes et al., 2014; Scott et al., 2011; Shemorry et al., 2013; Yang et al., 2013). Aberrant N-terminal acetylation can also lead to cellular dysfunction including drought-stress signaling in plants (Linster and Wirtz, 2018), human diseases (Dorfel and Lyon, 2015; Myklebust et al., 2015), and cancer (Kalvik and Arnesen, 2013).

Protein N-terminal acetylation is an irreversible modification that is catalyzed by a family of enzymes called N-terminal acetyltransferases (NATs). Dozens of NATs are present, varying in subunit composition, substrate specificity, and regulatory mechanisms (Deng and Marmorstein, 2021). Included amongst them are the three bacterial (RimI, RimJ, and RimL) (Pathak et al., 2016; Vetting et al., 2008; Vetting et al., 2005) homologues and a single archaeal NATs homologue (Liszczak and Marmorstein, 2013; Mackay et al., 2007); all of which tend to harbor relatively flexible N-terminal substrate specificity. Eukaryotes, however, possess more NATs (NatA-H) for more specialized activities and functions. Among them, five NATs (NatA, B, C, D, E) are conserved from yeast to human (Rathore et al., 2016; Starheim et al., 2012), associating with the ribosome to co-translationally acetylate nascent protein chains in a co-translational manner (Magin et al., 2017; Polevoda et al., 2008). NatA, B, C, and E are multi-subunit complexes, each consisting of at least one catalytic subunit and one auxiliary subunit, with the auxiliary subunits serving to modulate the activities of the respective catalytic subunits and act as an anchor to the ribosome, while the rest of the NATs function independently of auxiliary subunits. In addition to NatA, the NatE complex contains an additional catalytic subunit, Naa50, responsible for modifying NatE-type substrates (Deng et al., 2019; Deng et al., 2020a).

Substrate recognition by eukaryotic NATs is usually dictated by the first few residues of the substrate via the active site structure (Deng et al., 2019; Deng and Marmorstein, 2021; Deng et al., 2020b; Goris et al., 2018; Hong et al., 2017; Liszczak et al., 2013; Magin et al., 2015; Stove et al., 2016). The active site of NatA contains a relatively small binding pocket to accommodate a small, uncharged residue at position 1 such as alanine, valine or threonine residues that remains following initiator methionine (iMet)-cleavage (Liszczak et al., 2013). NatB has a hydrophobic pocket for a retained iMet and uses a histidine residue to recognize sequences with D/E/N/Q at position 2 via hydrogen bonding (Arnesen et al., 2009; Deng and Marmorstein, 2021). Interestingly, NatC and NatE share similar and relatively versatile substrate profiles toward substrates containing a retained iMet followed by residues other than D/E/N/Q (Arnesen et al., 2009; Van Damme et al., 2016). NatD is a highly selective enzyme with histones H4 and H2A as its only known substrates (Hole et al., 2011; Song et al., 2003), arising from extensive and specific interactions with each of the first four residues of its substrate (Magin et al., 2015). Although NATs vary in terms of their modes of substrate recognition, their mechanism of catalysis usually involves the use of dedicated residues to serve as general base and acid residues (Deng and Marmorstein, 2021).

The activity of some NATs is further regulated by other protein modulators. For example, within the human NatE complex, there is catalytic crosstalk between the two NAA10 and NAA50 catalytic subunits (Deng et al., 2019). Moreover, the activities of both NatA and NatE are inhibited to a different extent by a small protein binding partner called Huntingtin-interacting protein K (HYPK) (Arnesen et al., 2010; Deng et al., 2020a; Gottlieb and Marmorstein, 2018; Weyer et al., 2017). Inositol hexaphosphate (IP_6_, also named inositol hexakisphosphate or phytic acid), a small molecule, usually reserved for cellular signaling, appears to act as a stabilizing ligand for both NatA and NatE, by binding in a pocket between NAA15 and NAA10, and, thus, plays an indirect role in promoting NatA and NatE acetylation activity (Cheng et al., 2019; Deng et al., 2019; Deng et al., 2020a; Gottlieb and Marmorstein, 2018).

NatC is distinct from other NATs since it contains two eukaryotic conserved auxiliary subunits – Naa38 and Naa35 – that act together with the Naa30 catalytic subunit (Ochaya et al., 2019; Polevoda and Sherman, 2001; Starheim et al., 2009). Initial studies reported that NatC activity requires the interaction of all three subunits, since knock out of any single subunit produced a similar phenotype in yeast that was accompanied by diminished N-terminal acetylation of a cognate N-terminal substrate, MIRLK-(Polevoda and Sherman, 2001). However, it was subsequently reported that the Naa38 subunit is not required for *in vivo* acetylation of an Arl3p substrate with an N-terminal protein sequence of MPHLV-(Setty et al., 2004). In plants, knockout of *At*NAA35 did not lead to similar phenotypic effects as the deletion of *At*NAA30 (Pesaresi et al., 2003), and the single subunit of the *A. thaliana* analogue *At*NAA30 can rescue the knockout of yeast NatC complex subunits (Pesaresi et al., 2003). Similarly, recombinant human NAA30 (hMak3) was shown to have substrate-specific acetylation activity, even in the absence of its auxiliary subunits (Starheim et al., 2009). Thus, the functional roles of the NatC auxiliary subunits are not clear, particularly with respect to Naa38, which is not well-conserved across species. Although the recently reported crystal structures and associated biochemical studies of *S. cerevisiae* NatC (*Sc*NatC) by Grunwald *et al*. have provided some important insights into NatC substrate specificity and its catalytic mechanism (Grunwald et al., 2020), several questions remain unanswered. In particular, those regarding the mechanisms dictating the overlapping yet distinct substrate profiles of NatC and NatE and the influence of IP_6_ on NatC.

In this study, we report that yeast NatC complex formation of all three subunits is a prerequisite for normal NatC acetylation activity and that, like NatA and NatE, IP_6_ binds tightly to and stabilizes NatC. We also report the Cryo-EM structure of ~100 kDa *S. pombe* NatC (*Sp*NatC) bound to a NatE/C-type bisubstrate analogue and IP_6_, with related biochemistry, to reveal the molecular basis for IP_6_-mediated stabilization of the complex and the similar substrate profiles of NatE and NatC. Comparison of these studies with the recently published structural and functional studies of the *Sc*NatC complex (Grunwald et al., 2020) reveal evolutionarily conserved and divergent features of NatC.

## Results

### Naa38 is required for normal NatC acetylation activity in yeast

We found that overexpression of the *Sp*NatC complex containing full-length *Sp*Naa30 (residues 1–150), N-terminally truncated *Sp*Naa35 (residues 31–708, with deletion of divergent residues, 1-30) and *Sp*Naa38 (residues 48-116, with deletion of divergent residues, 1-47) in *E. coli* produced soluble protein that could be purified to homogeneity (**Figure 1a**). To evaluate the activity of the recombinant ternary *Sp*NatC, we used an *in-vitro* acetyltransferase activity assay with different peptide substrates. Consistent with previous studies, the recombinant *Sp*NatC is active toward both its own canonical substrate - “MLRF peptide” and the canonical NatE substrate – “MLGP peptide”, but with preference for the “MLRF peptide” (Grunwald et al., 2020) (**Figure 1b**, see methods section for full peptide sequences). To investigate the catalytic roles of Naa38, we could readily purify both binary (*Sc*Naa30^1-161^ and *Sc*Naa35^19-733^) and ternary *Sc*NatC complexes (*Sc*Naa30^1-161^ and *Sc*Naa35^19-733^, and *Sc*Naa38^1-70^) from *Saccharomyces cerevisiae* to compare their activities. Consistent with the studies by Grunwald *et al (Grunwald et al., 2020*), evaluation of the activities of the binary and ternary *Sc*NatC complexes toward the canonical NatC substrate (MLRF peptide) revealed that the ternary complex showed robust activity, while the binary complex showed compromised activity (**Figure 1b**). Thus, it appears that Naa38 is required for optimal NatC activity in yeast.

**Figure 1.**
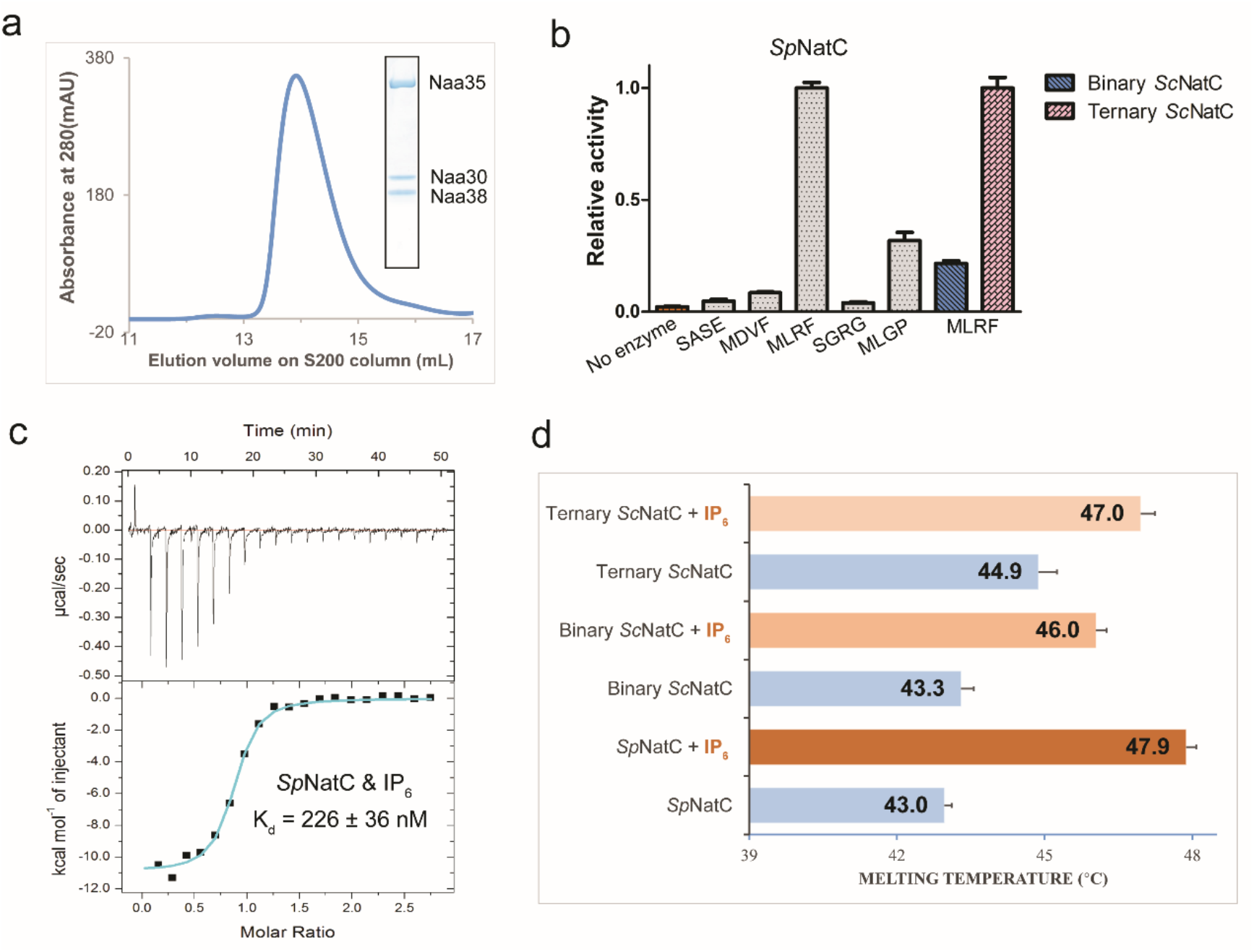
Binary NatC displays deficiency in acetylation activity and inositol hexaphosphate (IP_6_) binding stabilizes NatC complex formation. **(a)** Gel filtration elution profile of the ternary *Sp*NatC complex using a Superdex S200 column. Coomassie-stained SDS–PAGE of elution peak is shown in the box, with bands for corresponding *Sp*NatC subunits labeled. **(b)** Comparison of *Sp*NatC activity toward different peptide substrates. The activities shown in gray are normalized to the activity of *Sp*NatC toward MLRF peptide. The pink and blue columns show the activity of binary and ternary *Sc*NatC toward the MLRF peptide, respectively, and are normalized to the pink column. Errors are shown in SEM with n = 3. **(c)** Representative isothermal titration calorimetry (ITC) curve of IP_6_ titrated into *Sp*NatC. The ITC fitting information is N = 0.830 ± 0.00943 sites, ΔH = −1.091×10^4^ ± 177.0 cal mol^-1^, ΔS = −6.79 cal mol^-1^ deg^-1^. **(d)** Differential scanning fluorimetry assays of NatC alone or with IP_6_. Average calculated melting temperature transitions are indicated. Error bars in the figure indicate the SD of each sample, n = 3.

### Inositol hexaphosphate (IP_6_) binding contributes to yeast NatC complex stability

Earlier reports demonstrated that IP_6_ interacts with and stabilizes both the yeast and human NatA and NatE complexes (Cheng et al., 2019; Deng et al., 2019; Deng et al., 2020a; Gottlieb and Marmorstein, 2018), but does not appear to interact with human NatB (Deng et al., 2020b). To determine if IP_6_ could bind to the recombinant NatC complex, we employed isothermal titration calorimetry (ITC). We found that IP_6_ binds to *Sp*NatC with a Kd of ~225 nM (with a stoichiometry of ~1) (**Figure 1c**), similar to its Kd value for yeast NatA (Deng et al., 2019). Differential scanning fluorimetry (DSF) was used to evaluate a potential role of IP_6_ binding in NatC complex stability. We found that addition of IP_6_ increased the thermal stability of ternary *Sp*NatC, binary *Sc*NatC, and ternary *Sc*NatC, by ~4.9°C, 2.7°C, and 2.1°C, respectively. Thus, IP_6_ can provide additional thermal stability to the yeast NatC complexes (**Figure 1d**).

### NatC adopts a distinct NAT architecture

In order to understand the molecular details underlying the ternary NatC complex, its overlapping yet distinct substrate specificity with NatE, and the mode of IP_6_ stabilization, we performed single particle Cryo-EM analysis using ternary *Sp*NatC prepared in the presence of the CoA-Ac-MLGP bisubstrate conjugate and IP_6_. The MLGP sequence was selected for synthesis of the conjugate inhibitor as it is a predicted substrate for both NatC and NatE. A 3.16 Å-resolution Cryo-EM threedimensional (3D) map was determined from 607,131 particles, which were selected from 5,397 raw electron micrographs (**Supplementary Figure 1)**. Most areas of the EM map contained excellent sidechain density, with a local resolution of ~3 Å, suggesting the relative rigidity of the recombinant complex (**Supplementary Figure 1B and 2**). The atomic model was built *de novo* based on the EM map and details for the model refinement statistics can be found in **Table 1 (Supplementary Figure 2)**.

**Table 1.** Cryo-EM data collection, refinement and validation statistics.

|  |  |
| --- | --- |
|  | SpNatC with Bisubstrate and IP<sub>6</sub><br>EMD-23110<br>PDB: 7L1K |
| **Data collection and processing** |  |
| Magnification | 105,000 |
| Voltage (keV) | 300 |
| Electron exposure (e/Å<sup>2</sup>) | 42 |
| Defocus range (μm) | -1.0 to -3.0 |
| Pixel size (Å) | 0.84 |
| Symmetry imposed | C1 |
| Initial particles (no.) | 1,860,276 |
| Final particles (no.) | 607,131 |
| Map resolution (Å) | 3.16 |
| FSC threshold | 0.143 |
| Map resolution range (Å) | 2.5-3.5 |
| **Refinement** |  |
| Initial model used (PDB code) | - |
| Model resolution (Å) | 3.4 |
| FSC threshold | 0.5 |
| Model resolution range (Å) | - |
| Map sharpening *B* factor (Å<sup>2</sup>) | 159.1 |
| **Model composition** |  |
| Non-hydrogen atoms | 7216 |
| Protein residues | 883 |
| Ligands | 2 |
| ***B* factors (Å<sup>2</sup>)** |  |
| Protein | 26.97/110.48/71.91 |
| Ligand | 86.46/102.51/93.10 |
| **R.M.S. deviations** |  |
| Bonds lengths (Å) | 0.006 |
| Bond angles (°) | 1.036 |
| **Validation** |  |
| MolProbity score | 1.76 |
| Clash score | 4.10 |
| Poor rotamers (%) | 0.38 |
| **Ramachandran plot** |  |
| Favored (%) | 89.55 |
| Allowed (%) | 10.22 |
| Disallowed (%) | 0.23 |

The ternary *Sp*NatC complex contains three proteins, the catalytic Naa30 subunit, the large auxiliary Naa35 subunit, and the small auxiliary Naa38 subunit. Upon superficial inspection of the complex, it appears as though it is only formed by two proteins, as Naa38 is embedded within the Naa35 fold to form a “composite” auxiliary subunit (**Figure 2a and 2b)**. While Naa30 displays a canonical NAT fold with mixed α/β secondary structure, the Naa35/38 complex forms a clamp that pinches Naa30 on opposite sides. This clamp-like structure is distinct from the architecture of the large auxiliary subunits of NatA and NatB, which surround the base of their respective catalytic subunits (Deng et al., 2020b; Gottlieb and Marmorstein, 2018; Hong et al., 2017; Liszczak et al., 2013). The Naa35 architecture is mostly helical, with 25 helices, three short β-strands in its N-terminal region and a single β-hairpin at its C-terminal region that are unique to the NatC complex (**Figure 2b and Supplementary Figure 3**). Naa38 adopts an Sm-like fold, with an N-terminal α-helix, followed by a sharply bent β-sheet consisting of 5 five antiparallel β-strands (**Figure 2b and Supplementary Figure 4)**. The Naa35/Naa38 clamp contains α4-α13 from Naa35 at its base, and α14-α25 and the C-terminal β-hairpin of Naa35 flanking one side of Naa30. The opposing side of Naa30 is flanked by a composite of the N-terminal end of Naa35 (α2, α3, β1-β3) and the entirety of Naa38 (**Figure 2a and 2b**). Consistent with the composite nature of the Naa35-Naa38 interaction, the buried surface area between Naa38 and Naa35 is ~1925 Å^2^, which is notably higher than the area buried between Naa30 and Naa35 (~1735 Å^2^), and between Naa30 and Naa38 (~197 Å^2^).

**Figure 2.**
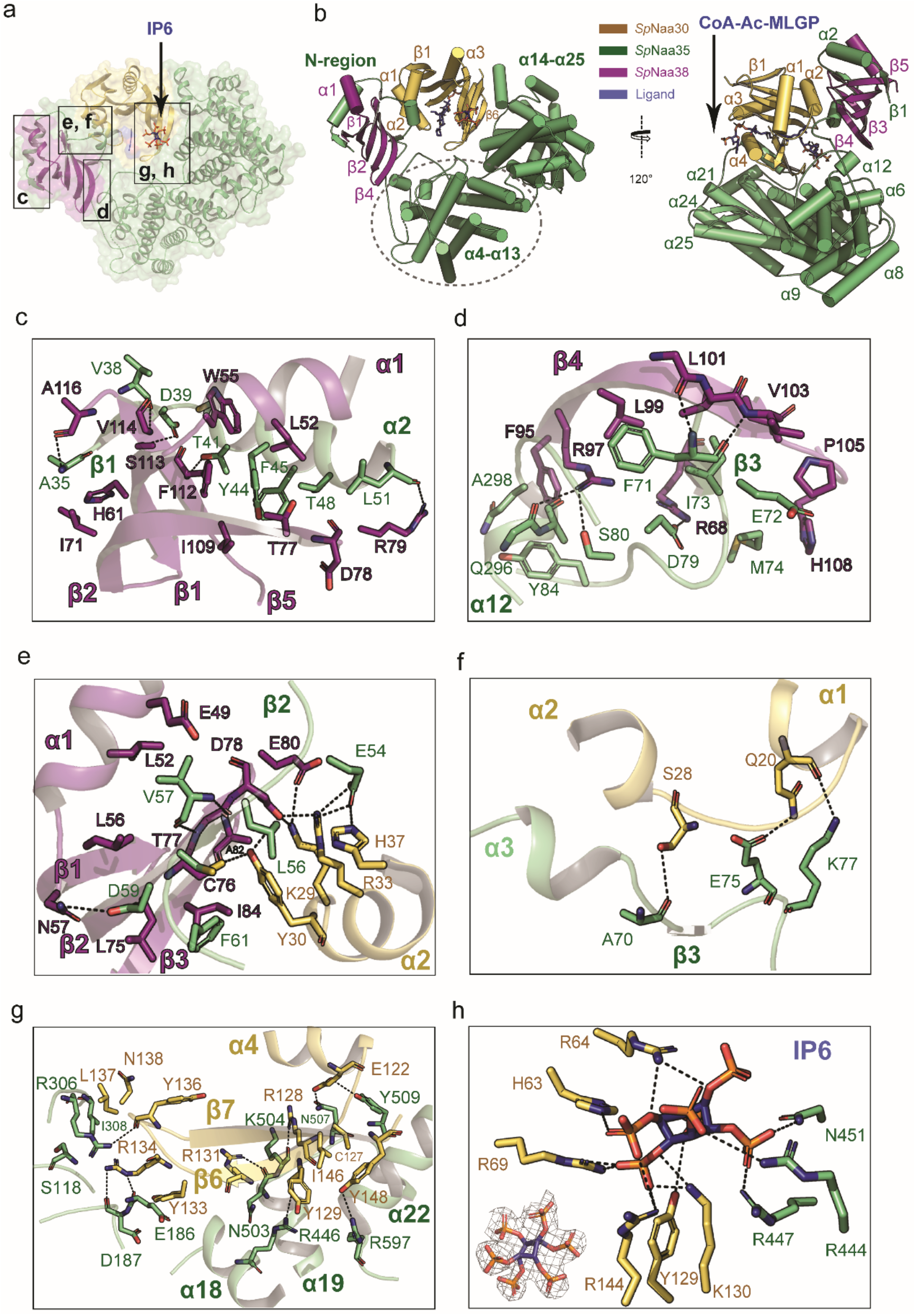
Overall structure of *Sp*NatC reveals intimate interactions among all subunits. **(a**) Naa30 (bright orange), Naa38 (dark purple) and Naa35 (lime green) are shown in transparent surface and cartoon, with the boxed area labeled. **(b)** The CoA-MLGP bisubstrate conjugate and inositol hexaphosphate (IP_6_) are shown in sticks and colored in blue. **(c)** Zoom-in view of the first sub-interface between Naa38 and Naa35. Hydrogen bonds are indicated by dashed black lines. For simplicity, only some of the hydrophobic interactions are shown. **(d)** Zoom-in view of the second sub-interface between Naa38 and Naa35. **(e)** Zoom-in view of the third sub-interface between Naa38 and Naa35, which also involves some residues from Naa30 (light orange). **(f)** Zoom-in view of the interface between Naa35 and Naa30 α1-α2 segment. **(g)** Zoom-in view of the interface between Naa35 and Naa30 β6-β7 segment. **(h)** Zoom-in view depicting key residues involved in interactions with IP_6_. The lower left corner shows the fit of the IP_6_ molecule in the EM density map. The contour level is 5.0 sigma.

### Naa38 plays a key role in the NatC complex

The structure of *Sp*NatC was consistent with our biochemical data suggesting that Naa38 plays important structural and functional roles within the NatC complex. A more detailed view of the Naa35-Naa38 interface reveals that it is nucleated by the formation of three anti-parallel β-sheets between Naa35 and Naa38, resulting in extensive hydrogen bonding and van der Waals interactions, which also involve the α2 and α3 helices of Naa35. These interactions can roughly be divided into three regions according to the relative position of the β-sheets (**Figure 2**).

The first sub-interface involves the Naa35 β1-α2 segment (**Figure 2c**). Hydrogen bonds are formed between the backbone nitrogen of Naa35-Ala35 and the backbone carbonyl of Naa38-Ala116, the backbone carbonyl of Naa35-Val38 and the backbone nitrogen of Naa38-Val114, the sidechains of Naa35-Asp39 and of Naa38-Ser113, the sidechain of Naa35-Thr41 and the backbone nitrogen of Naa38-Phe112, and the backbone carbonyl of Naa35-Leu51 and the sidechain of Naa38-Arg79. This region also features a large area of van der Waals interactions involving residues Ala35, Gly36, Tyr37, Tyr44, Phe45, Ala47, and Thr48 from Naa35 and residues Gly48, Leu52, Trp55, His 61, Ile71, Thr77, Asp78, Ile109, Arg115, and Ala116 from Naa38.

The second sub-interface is mediated by Naa35-β3 and α12 (**Figure 2d**). Hydrogen bonds are formed between the backbone nitrogen of Naa35-Phe71 and the backbone carbonyl of Naa38-Leu101, the backbone carbonyl of Naa35-Phe71 and the backbone nitrogen of Naa38-Leu103, the sidechains of Naa35-Asp79 and Naa38-Arg68, the sidechains of Naa35-Ser80 and Naa38-Arg97, and the backbone carbonyl of Naa35-Gln296 and the sidechain of Naa38-Arg97. Residues involved in van der Waals interactions include Ala70, Glu72, Ile73, Met74, Tyr84, Ala 297, Gln 298, Val30, and Ala301, from Naa35, and residues Phe95, Ala98, Leu99, Val102, Ile104, Pro105, and His108 from Naa38.

The third sub-interface is mediated by Naa35-β2 but also involves extensive interactions with the catalytic Naa30 subunit (**Figure 2e**). The hydrogen bonds between Naa35 and Naa38 are formed between the sidechains of Naa35-Asp59 and Naa38-Asn57, the backbone nitrogen of Naa35-Val57 and the backbone carbonyl of Naa38-Thr77, and the backbone carbonyl of Naa35-Val57 and the backbone nitrogen of Naa38-Thr77. Naa38 engages with the α2 helix of the Naa30 subunit to form an extensive hydrogen bonding network via the Naa30-Tyr30 sidechain to the backbone carbonyl of Naa38-Ala82, and the sidechains of Naa38-Cys76 and Asp78; the sidechain of Naa30-Lys29 hydrogen bonds to the sidechains of Naa38-Asp78 and Glu80; and between the sidechains of Naa30-Arg33 and Naa38-Asp78. As the α1-α2 loop of Naa30 plays a key role in protein N-terminal substrate recognition, these Naa38-Naa30 interactions likely play a key role in substrate recognition by NatC (see discussion). Finally, the Naa30-α2 helix also interacts with Naa35 where Naa35-Glu54 forms hydrogen bonds with Naa30-Arg33 and His37. Within this sub-interface, the hydrophobic interactions are primarily mediated by residues Glu49, Leu52, Leu56, Leu75, Cys76, Asp78, and Ile84 from Naa38 and residues Leu56, Cys58, Asp59 and Phe62 from Naa35. Taken together, it appears that Naa38 plays key structural and, likely, functional roles in NatC.

### The substrate binding α1-α2 loop and β6-β7 segments of Naa30 are buried within Naa35

Naa30 uses its α1-α2 loop (**Figure 2f)** and β6-β7 segments to interact with Naa35 **(Figure 2g)**. Analogous segments in the NatA and NatB complexes are used for N-terminal substrate recognition. In these complexes, however, the β6-β7 segment is exposed to solvent (Deng et al., 2020b; Gottlieb and Marmorstein, 2018; Hong et al., 2017; Liszczak et al., 2013). By contrast, this interaction between Naa30 and Naa35 is facilitated by hydrogen bonds between the Naa30 α1-α2 loop and Naa35 α3-β3 segment: the sidechain of Naa30-Ser28 and the backbone carbonyl of Naa35-Ala70; between the sidechains of Naa30-Gln20 and Naa35-Glu75; and the backbone carbonyl of Naa30-Gln20 and the sidechain of Naa35-Lys77.

Notably, the interactions between Naa30 β6-β7 and Naa35 are significantly more extensive than those found in the NatA and NatB complexes **(Figure 2g)**. Here, hydrogen bonds are formed between the sidechain of Naa30-Glu122 and the sidechains of Naa35-Gln507 and Tyr509, the sidechain of Naa30-Arg128 and the backbone carbonyl of Naa35-Lys504, the backbone carbonyl of Naa30-Tyr129 and the sidechain of Naa35-Arg446, the sidechain of Naa30-Arg131 and the backbone carbonyl of Naa35-Gln503, the sidechain of Naa30-Arg134 and the backbone carbonyls of both Naa35-Glu186 and Asp187, the backbone carbonyl of Naa30-Tyr136 and the sidechain of Naa35-Arg306, and the sidechains of Naa30-Tyr148 and Naa35-Arg597. Van der Waals interactions are observed between residues Cys127, Leu132, Tyr133, Leu137, Asn138, Phe143, Ile146, and Tyr148 from Naa30, and residues Ile308, Asn441, Cys443, Leu505, Phe581, Ser590, Tyr591, and Ala594 from Naa35. Together, it appears that the Naa35 auxiliary subunit makes extensive contacts with the protein N-terminal substrate binding loops of Naa30 to influence substrate recognition.

### IP_6_ is bound at the interface between Naa30 and Naa35 in close proximity to the peptide substrate binding site

EM density corresponding to a bound IP_6_ molecule is well-resolved, revealing a IP_6_ binding pocket at the interface between the Naa30 and Naa35 subunits (**Figure 2h)**. Notably, the IP_6_ binding region is distinct from its binding sites in the NatA and NatE complexes (Deng et al., 2019; Deng et al., 2020a; Gottlieb and Marmorstein, 2018). In *Sp*NatC, IP_6_ engages in polar interactions with Naa30 residues His63, Arg64, Arg69, Tyr129, Lys130, and Arg144, and Naa35 residues Arg444, Arg447, and Asn451. Interestingly, although the IP_6_ binding pocket of NatC features a similar chemical environment to the one found in the NatA and NatE complexes, the location in the architecture of these NAT complexes differs. In NatC, IP_6_ is located close (~8 Å) to the peptide substrate entrance site, while the pocket found in NatA and NatE is close to the Ac-CoA entrance site (**Figure 3)**. Therefore, the NatC-bound IP_6_ molecule could be in position to alter NatC activity. To evaluate this possibility, we assayed the influence of IP_6_ on NatC activity and found that incubation of the NatC complex IP_6_ produced a relatively modest inhibitory effect and had a inhibitory effect on *Sp*NatA and hNatB, (**Supplementary Figure 5**). While we cannot exclude the possibility that this inhibitory effect may be an artifact of the radioactive assay, it appears that IP_6_ plays a predominantly structural role in NatC.

**Figure 3.**
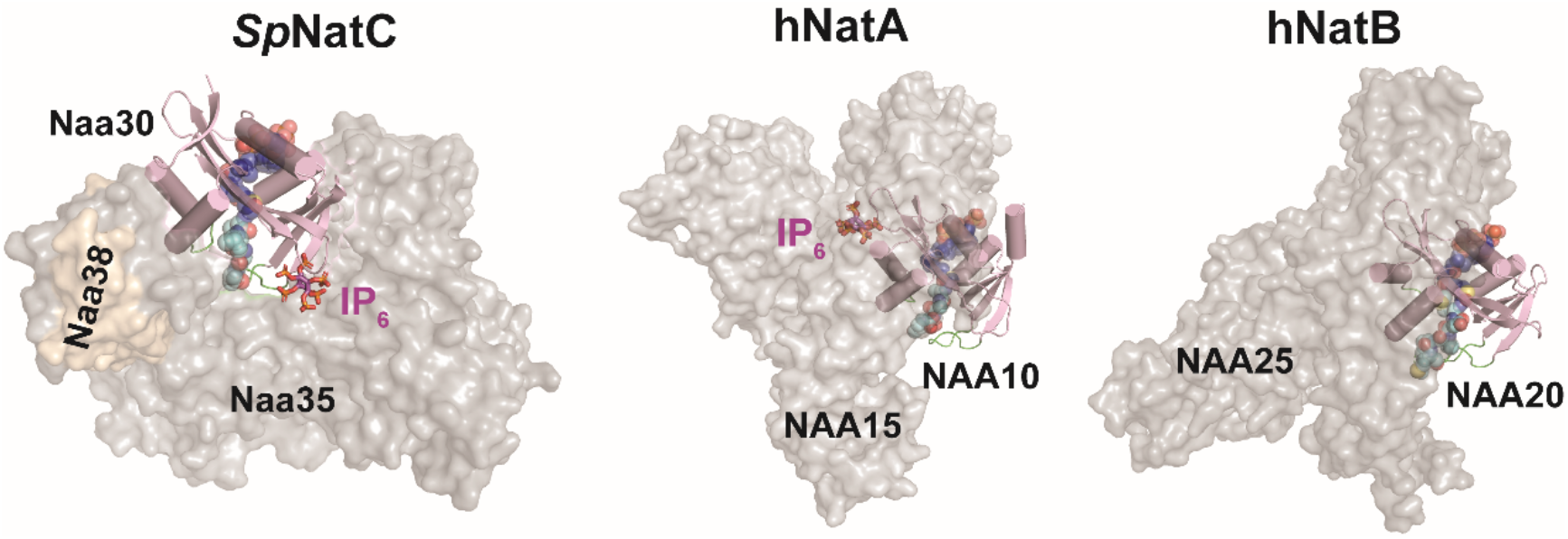
Comparison of NatC with NatA and NatB reveals divergent NatC architecture. The auxiliary subunits are shown as transparent surface in grey or brown, while the catalytic subunits are shown as cartoon in light pink (catalytic subunits are aligned in the same orientation). The blue spheres, cyan spheres, and magenta sticks represent Ac-CoA, peptide substrate, and IP_6_, respectively. The highlighted green loops shown are α1-α2 and β6-β7 substrate binding loops of the catalytic subunits. The PDB models for generating this figure: *Hs*NatA with IP_6_, PDB: 6C9M (the bisubstrate analogue shown is aligned from *Sp*NatA, PDB:4KVM); *Hs*NatB with the bisubstrate analogue, PDB: 6VP9; model of *Sp*NatC with the bisubstrate analogue and IP_6_, PDB: 7L1K. As NatE is a complex with dual catalytic subunits and shares the same auxiliary subunit with NatA, for simplicity, it is not compared in this figure.

### NatC architecture displays significant differences from NatA, NatE and NatB

Overall, there are three notable differences between NatC and the other multi-subunit NATs – NatA, NatE and NatB. First, the auxiliary subunits of NatA, NatE and NatB contain only helical secondary structure, forming 10-13 conserved tetratricopeptide repeat (TPR) motifs (Deng et al., 2019; Deng et al., 2020a; Deng et al., 2020b; Gottlieb and Marmorstein, 2018; Hong et al., 2017; Liszczak et al., 2013). In contrast, the NatC auxiliary subunits (Naa35 and Naa38) do not contain TPR repeats and but do contain several beta strands that make key interactions in the complex (**Figure 2b)**. Secondly, the auxiliary subunits of NatA, NatE, and NatB form a closed ring-like cradle to completely wrap around their corresponding catalytic subunit. In contrast, NatC auxiliary subunits arrange themselves into a clamp-like structure to only wrap roughly half-way around the Naa30 catalytic subunit (**Figure 3)**. Thirdly, the Naa30 β6-β7 peptide substrate recognition loop is buried within the auxiliary subunit, while the corresponding loops in NatA, NatE, and NatB are largely exposed to solvent (**Figure 3)**. Thus, the relative orientation of the catalytic subunit to the auxiliary subunit (s) is different in NatC relative to NatA and NatB. In turn, this has implications in altering the NatC substrate binding mode: the NatC substrate binding tunnel is roughly perpendicular to the Naa30-Naa35 interface, while the substrate binding tunnel of NatA, NatE, and NatB is parallel to the catalytic-auxiliary subunit-interface (**Figure 3)**. Taken together, the NatC complex forms an architecture that is distinct from other multi-subunit NATs.

### Substrate recognition by Naa30 shows similarity to NAA50

Overall, Naa30 displays a typical NAT fold containing four α-helices and seven β-strands with similar substrate binding modes: Ac-CoA enters the catalytic active site through a groove formed by α3 and α4 of the catalytic subunit, while the peptide substrate enters the active site on the opposite side of the catalytic subunit flanked by the α1–α2 and β6-β7 peptide substrate binding loops (Deng and Marmorstein, 2021) (**Figure 4a and 4b)**. In the Cryo-EM map, density for the CoA-Ac-MLGP conjugate bisubstrate is well-resolved, allowing us to confidently model both the CoA portion and all four residues of the peptide portion (**Figure 4c)**.

**Figure 4.**
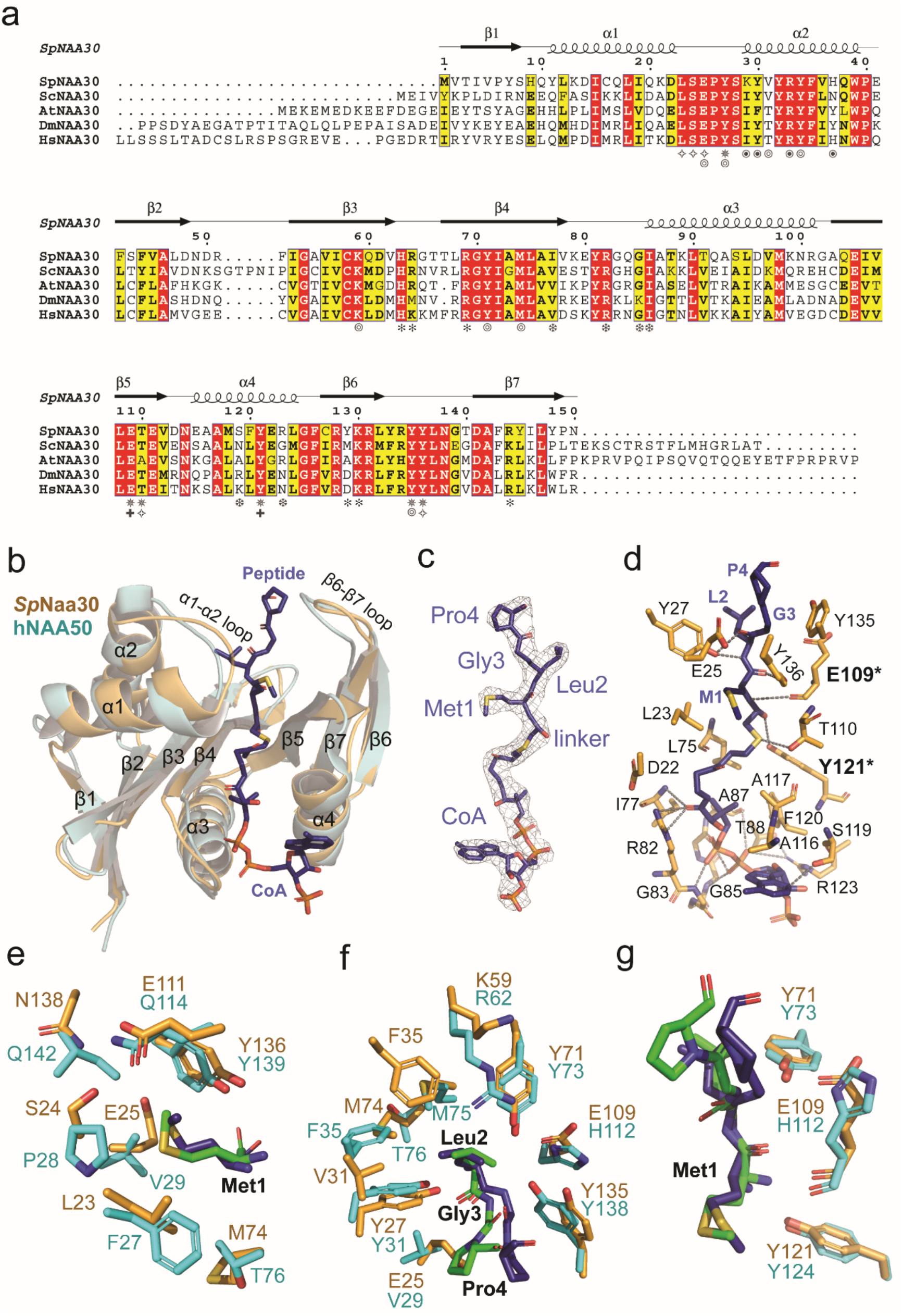
Substrate recognition by Naa30 is similar to NAA50. **(a)** Sequence alignment of Naa30 orthologs from *S. pombe* (Sp), *S. cerevisiae* (Sc), *A. thaliana* (At), *D. melanogaster* (Dm), and *H. sapiens* (Hs). Numbering and secondary structure elements for *Sp*Naa30 are indicated above the sequence alignment. Residues of *Sp*Naa30 that contact the peptide backbone (*), CoA (Φ), Met1 (✧)x, Leu2 (⦾), Naa38 (⦿), IP_6_ (⦿), and catalytic residues (+) are indicated. **(b)** Structural alignment of *Sp*Naa30 (bright orange) with *Hs*NAA50 (cyan), with secondary structure elements indicated. **(C)** The fit of the bisubstrate inhibitor in the EM density map. The contour level is 5.0 sigma. **(d)** Highlighted polar and hydrophobic interactions between CoA-Ac-MLGP and *Sp*Naa30 are depicted in the 3D view. **(e)** Residues forming a hydrophobic pocket surrounding the substrate peptide Met1 sidechain are shown in sticks (Orange, *Sp*Naa30; Cyan, *Hs*NAA50) **(f)** Residues forming a hydrophobic pocket surrounding the substrate peptide Leu2 sidechain are shown in sticks. (Orange, *Sp*Naa30; Cyan, *Hs*NAA50) **(g)**. Residues proposed as catalytic residues are shown in sticks. (Orange, *Sp*Naa30 E109 and Y121; Cyan, *Hs*NAA50 Y73 and H112).

Naa30 harbors a conserved Ac-CoA binding motif R_82_XXG_85_XA_87_ where Ac-CoA binding is mediated by a series of hydrogen bonds, mainly to the pyrophosphate group. Specifically, the sidechains of Arg82, Ser119, Arg123 and backbone atoms from Ile77, Gly85, Ile86, and Ala87 contribute to Ac-CoA hydrogen bonding (**Figure 4d)**.

In our NatC model, Naa30 binds the peptide substrate mainly through peptide backbone hydrogen bonding and hydrophobic pockets for the sidechains of the first two peptide residues. Direct hydrogen bonds are formed between the backbone amide groups of the peptide residues 1 and 2 to Naa30 residues Tyr27, Glu109, Thr110, Tyr121, and Tyr136 (**Figure 4d)**. The binding pocket for the peptide Met1 is surrounded by residues Leu23, Ser24, Glu25, Glu111, and Tyr136 (**Figure 4e)**, while the binding pocket for peptide Leu2 is half-open and surrounded by Tyr27, Val31, Tyr34, Phe35, Lys59, Tyr71, and Met74 (**Figure 4f)**. Beyond the first two residues, there are no direct contacts between the peptide substrate and Naa30, with the exception of distant contacts between the backbones of peptide Gly3 and peptide Pro4 with Naa30-Tyr135 (4-4.5 Å) and Naa30-Glu25 (~4 Å), respectively. These two Naa30 residues likely play a role in recognizing positions 3 and 4 of high-affinity NatC peptide substrates, as supported by the recent report by Grunwald *et al*. (Grunwald et al., 2020)

Previous studies have described the overlapping substrate profile of NatC and NAA50. Since our NatC model contains the canonical NatE (NAA50)-type peptide substrate (Met2-Leu2-Gly3-Pro4) in the bisubstrate inhibitor, we were able to evaluate this relationship by visual inspection of the inhibitor-bound *Sp*Naa30 subunit from the model. The *Sp*Naa30 model aligns well with the *Hs*NAA50 structure (PDB: 3TFY) bound to the same MLGP peptide fragment, with a root-mean-square deviation (RMSD) of 1.23 Å (over 105 common C_α_ atoms) (**Figure 4b**). A more detailed view of the peptide substrate binding site reveals that Met1 sits in similar binding pockets in both structures (**Figure 4e)**, consistent with their high specificity for Met1. Specificity for Leu at position 2 is also similar. Notably, residues NatC-Tyr27 and Tyr71, which are responsible for Leu2 backbone recognition align well with the corresponding residues in NAA50 (**Figure 4f)**. For Leu2 sidechain recognition, the residues, Naa30-Val31 and Lys59 are replaced by bulkier residues, NAA50-Phe35 and Arg62 (**Figure 4f)**, respectively. However, Naa30-Phe35, Ala73 and Met74 are replaced by NAA50-Phe35, Met75 and Thr76 (**Figure 4f),** perhaps explaining the greater tolerance of NAA50 for other residues at position 2. In both models, the peptide substrates begin to diverge from the architecture of their respective models at residue 3, where the Cα atoms of the corresponding peptide Gly3 residues of both models are ~3 Å apart. This also seems to be correlated with a shift in the *p*-hydroxyl group of a nearby conserved Tyr residue (Naa30-Tyr135 and NAA50-Tyr138) by about 2.5 Å (**Figure 4g**). The corresponding peptide Pro4 residues also point in different directions (**Figure 4g**). We propose that these differences in peptide substrate positioning beyond residue 2 is mediated by the different active site environments of Naa30 and NAA50 in this region, thus allowing these two enzymes to harbor varying activities toward substrates with the same residues at positions 1 and 2 but differing residues at position 3 and beyond (See discussion). Taken together, this comparison explains the overlapping yet distinct substrate profiles of NatC and NatE.

### Mutational analysis reveals that Naa30 Glu-109 and Tyr-121 play key catalytic roles

In order to evaluate key Naa30 residues identified in our model and their functional roles in substrate binding and catalysis, we used a radioactive *in vitro* acetyltransferase activity assay in conjunction with the MLRF peptide as the peptide substrate to kinetically characterize WT and mutant *Sp*NatC proteins. Each mutant was purified to homogeneity and displayed identical gel filtration chromatography elution profiles (data not shown), consistent with their native folding and complex formation. Consistent with our structural observations, mutation of a majority of residues involved in peptide substrate recognition resulted in an increase in K_m_ (**Table 2**). As the “YY motif” is shown to be conserved in most NATs (Stove et al., 2016) (except NatD/NAA40), single mutation of either Naa30-Tyr135 or Tyr136 to an alanine resulted in a significant loss in activity, with the Y136A mutant displaying almost no detectable activity. In previous studies, NAA50 residues Tyr73 and His112 were proposed to contribute to catalysis (Liszczak et al., 2011). The corresponding residues in Naa30, Tyr71 and Glu109, however, had an unexpected effect on catalysis. Instead, Naa30-Y71A or Y71F primarily altered the Km, which ultimately resulted in a minor decrease (12-67%) in catalytic efficiency in comparison with WT. On the other hand, mutation of Naa30-Glu109 to either alanine or glutamine led to >90% loss of catalytic efficiency, impacting both K_m_ and k_cat_. NAA50-Tyr124 and hNatB-Tyr123 were recently proposed to play analogous catalytic roles (Deng et al., 2020b), and the corresponding mutation of Naa30-Tyr121 to alanine demonstrated that NatC activity was almost completely abolished (**Table 2**). While Glu109 is positioned to function as a general base to deprotonate the α-amino group of the peptide substrate, Tyr121 is positioned to function as a general acid to re-protonate the CoA leaving group. This is consistent with the proposed roles of the analogous Glu118 and Tyr130 residues in *Sc*NatC *(Grunwald et al., 2020)*, as well as the conserved nature of these residues (**Figure 4a**). In addition, Naa30-Asn114 is highly conserved among all NATs (data not shown) and it was recently suggested to play a structural role in orienting the catalytic tyrosine residues (*Sp*Naa30-Tyr135 and Tyr136) by ensuring the proper position of the active site helix (α4) for interaction with Ac-CoA (Deng et al., 2020b). Consistent with the importance of Asn114, we observed that Naa30-N114A exhibited an ~50% loss in activity. Taken together, the conserved Naa30-Glu109 and Tyr121 residues likely play key catalytic roles in NatC complex activity (**Figure 4g)**.

**Table 2.** *Sp*NatC complex (WT and mutant *Sp*Naa30) catalytic parameters.

| Substrate | Protein<br>(*SpNatC*) | $k_{cat}^*$<br>($\text{min}^{-1}$) | Relative $k_{cat}$<br>(normalized<br>to WT) | $K_m^*$<br>($\mu\text{M}$) | Relative $K_m$<br>(normalized<br>to WT) | $k_{cat}/K_m$<br>(normalized<br>to WT) |
| --- | --- | --- | --- | --- | --- | --- |
| *gag*<br>Peptide<br>(MLRF-) | WT | $23.9 \pm 1.4$ | 1.0 | $7.6 \pm 2.1$ | 1.0 | 1.0 |
| | S24A | $22.7 \pm 1.6$ | 0.95 | $10.3 \pm 3.2$ | 1.4 | 0.70 |
| | E25A | $49.3 \pm 2.3$ | 2.1 | $19.7 \pm 3.6$ | 2.6 | 0.79 |
| | Y27A | $7.33 \pm 0.5$ | 0.31 | $3.0 \pm 1.0$ | 0.39 | 0.79 |
| | Y71A | $44.7 \pm 2.8$ | 1.9 | $16.1 \pm 4.2$ | 2.1 | 0.88 |
| | Y71F | $30.2 \pm 2.3$ | 1.3 | $29.6 \pm 8.5$ | 3.9 | 0.32 |
| | M74A | $13.7 \pm 1.1$ | 0.57 | $18.7 \pm 6.1$ | 2.5 | 0.23 |
| | E109A | $10.5 \pm 1.6$ | 0.44 | $28.7 \pm 16$ | 3.8 | 0.11 |
| | E109Q | $3.2 \pm 0.9$ | 0.13 | $201 \pm 130$ | 27 | 0.0050 |
| | N114A | $9.5 \pm 0.6$ | 0.40 | $5.6 \pm 1.7$ | 0.74 | 0.53 |
| | Y121A | $1.2 \pm 0.1$ | 0.050 | $16.0 \pm 5.4$ | 2.1 | 0.023 |
| | Y121F | $1.4 \pm 0.2$ | 0.060 | $7.9 \pm 4.3$ | 1.0 | 0.057 |
| | Y135A | $6.3 \pm 0.8$ | 0.26 | $9.5 \pm 5.7$ | 1.3 | 0.21 |
|  | Y136A | N.A | N.A | N.A | N.A | N.A |
| *values indicated represent mean ± S.D. of best-fit curves | | | | | | |

## Discussion

Here, were report that the notably small *Sp*Naa38 auxiliary subunit cooperates with the large Naa35 auxiliary subunit to stabilize the active N-terminal acetyltransferase NatC complex for its robust N-terminal acetylation activity. Through structure/function studies, we observed that Naa38 interacts with Naa30-α2, a region that directly contributes to recognition of peptide substrate backbone and sidechain residues. Based on these observations and similar *Sc*NatC complex biochemical and structural findings described by Grunwald *et al. (Grunwald et al., 2020)*, it appears that Naa38 is essential for normal NatC acetylation activity. Therefore, we propose that Naa38 plays a conserved role in yeast. Future studies are needed to further interrogate this in the plant or human systems, as some reports suggest that Naa38 may not be as important in higher eukaryotes (Pesaresi et al., 2003; Starheim et al., 2009), but there has also been a report of a human developmental disorder linked to *NAA38* deletion (Zhao et al., 2016).

In *S. pombe*, we have found that the ternary *Sp*NatC complex displays acetylation activity toward both canonical NatC and NatE-type peptide substates, but with a preference for NatC-type substrates. While recognition of residues 1 and 2 of the cognate substrate appears highly homologous between the Naa30 and NAA50 catalytic subunits of NatC and NatE, respectively, the substrate binding pockets responsible for recognizing substrate residues beyond the penultimate sidechain diverge significantly. This likely explains the divergence in substrate profiles observed between these two NATs.

IP_6_ was previously found to play a stabilizing role at the interface between the NAA15 and NAA10 subunits of the NatA and NatE complexes (Deng et al., 2019; Deng et al., 2020a; Gottlieb and Marmorstein, 2018), (Cheng et al., 2019). Here, we find that IP_6_ also appears to play a stabilizing role in both *Sc*NatC and *Sp*NatC, and that the *Sp*NatC structure reveals that IP_6_ binds at the Naa35-Naa30 interface located near the peptide substrate entrance site, although it does not appear to play a major direct role in modulating NatC activity.

Grunwald *et al*. recently reported X-ray crystal structures of *Sc*NatC in several liganded states along with associated biochemical studies *(Grunwald et al., 2020)*. Overall, their findings are in agreement with our findings on *Sp*NatC reported here. There are, however, some notable differences when comparing NatC from *S. pombe* and *S. cerevisiae*, which have implications for the conserved and unique features of NatC.

The *Sc*NatC structures bound to cognate NatC-type peptide substrates shows specificity for the first four residues, which is consistent with their reported peptide substrate mutation and binding data. In contrast, our structure, featuring *Sp*NatC bound to a peptide sequence that is optimal for NatE (NAA50) acetylation, shows *Sp*Naa30’s specificity for only the first two residues. This mode of recognition is consistent with recognition for these same residues by NAA50, while there is significant divergence in the regions of the peptide binding grooves of Naa30 and NAA50 that are responsible for interaction with substrates residues beyond residue 2. This comparison suggests that discrimination between Naa30 and NAA50 substrates is largely dictated by the identities of the third and fourth residues in an N-terminal substrate. As NatC-type and NatE-type substrates can include many different hydrophobic/amphipathic N-termini (ML- MI-, MF, MW-, MV-, MM-, MH-, and MK-) (Ree et al., 2018), it may be difficult for a single enzyme to cover acetylation of such a broad repertoire of protein N-termini. Given that Naa50 is inactive in yeast (Deng et al., 2019), we propose that the overlapping substrate profile for residues at positions 1 and 2 by NatC and NatE may have evolved in higher eukaryotic cells to fully cover the acetylation on these types of N-terminal substrates.

Grunwald *et al*. identified several electropositive regions (EPR) on *Sc*NatC that are implicated in ribosome association. Of these regions, labeled EPR1-EPR4 (Grunwald et al., 2020) (**Figure 5**), mutation of only EPR2 was found to influence ribosome association. Comparison with the electrostatic surface potential of *Sp*NatC shows notable differences with *Sc*NatC (**Figure 5)**. Specifically, EPR2 is not present in *Sp*NatC, since the *Sc*NatC “*Sc*Naa35 tip” is missing in *Sp*NatC. While EPR1 is conserved, this electropositive patch harbors the IP_6_-binding pocket in *Sp*NatC and, therefore, is likely the IP_6_ binding site in *Sc*NatC. In addition, the surface region of *Sp*NatC corresponding to EPR3 is broader compared to the corresponding surface of *Sc*NatC and is located close to where *Sc*NatC-EPR2 would be. It is therefore possible that *Sp*NatC utilizes EPR3 to compensate for the absence of EPR2. Taken together, this comparison suggests that the mode of ribosome association utilized by *Sp*NatC and *Sc*NatC and, by extension, NatC complexes from other species, may differ. Future studies would investigate the molecular details of these differing interactions.

**Figure 5.**
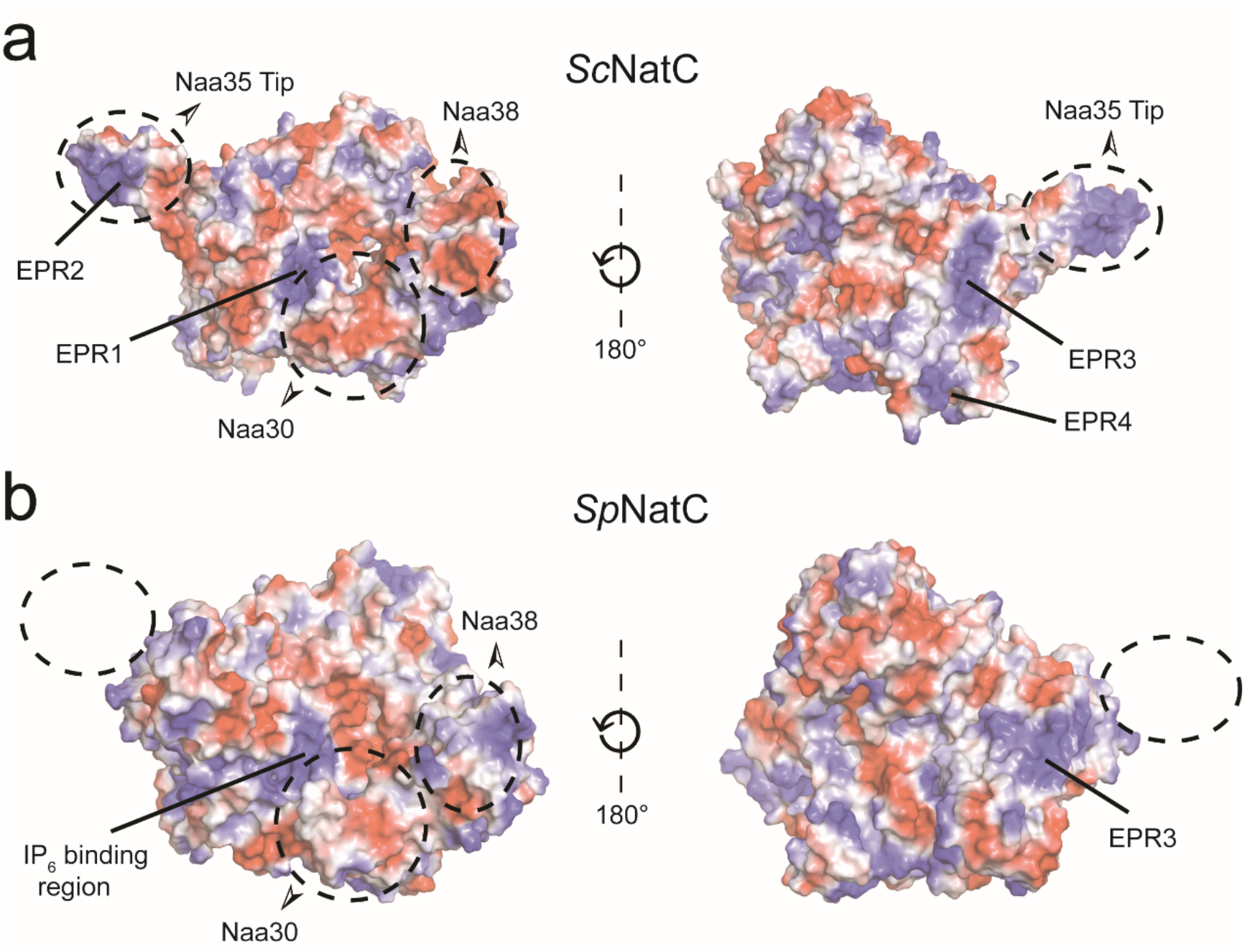
Divergent structural features exist between NatC in *S. pombe* and *S. cerevisiae*. **(a)** Electrostatic surface potential of the *Sc*NatC structure (PDB: 6YGB) with EPRs indicated. **(b)** Electrostatic surface potential of *Sp*NatC (PDB: 7L1K) aligned with the orientations shown in (a).

NatC has been shown to be important for proper chloroplast (Pesaresi et al., 2003) and mitochondrial (Van Damme et al., 2016) function, essential for embryonic development (Wenzlau et al., 2006), cell viability and p53-dependent apoptosis (Starheim et al., 2009), and as a potential therapeutic target in cancer (Mughal et al., 2015). In light of these important connections to NatC function, the studies reported here have important implications for health and disease.

## Materials and Methods

### *Sp*NatC Complex Expression and Purification

*Sp*Naa35^31-708^ (with truncation of the N-terminal 30 residues) and full-length *Sp*Naa30^FL^ were cloned into a modified pET DUET vector containing an N-terminal His_6_-SUMO tag. *Sp*Naa38^48-116^ (with truncation of the N-terminal 47 residues) was cloned into a pCDF vector with an N-terminal His6 tag. To obtain the heterotrimeric NatC complex (WT ternary *Sp*NatC), these two plasmids were used to co-transform Rosetta (DE3)pLysS competent *E. coli* cells, which was then cultured to grow at 37 °C in the presence of both ampicillin (100 μg/mL) and streptomycin (50 μg/mL). When the absorbance OD_600_ reached ~ 0.7, protein expression was induced at 16°C with 0.5 mM isopropyl 1-thio-β-galactopyranoside (IPTG) overnight. Cells were harvested by centrifugation, resuspended, and lysed by sonication in lysis buffer containing 25 mM Tris, pH 8.0, 300 mM NaCl, 10 mg/ml PMSF (phenylmethanesulfonylfluoride). The lysate was clarified by centrifugation and passed over Nickel-NTA resin (Thermo Scientific), which was subsequently washed with ~10 column volumes of wash buffer containing 25 mM Tris, pH 8.0, 300 mM NaCl, 20 mM imidazole, 10 mM 2-mercaptoethanol (βME). The protein was eluted in batches with buffer containing 25 mM Tris, pH 8.0, 300 mM NaCl, 200 mM imidazole, 10 mM βME. After elution, His_6_-tagged Ulp1 protease was added to the eluate to cleave the His_6_-SUMO tag and dialyzed further into buffer containing 25 mM sodium citrate monobasic, pH 5.5, 10 mM NaCl and 10 mM βME. Protein was purified with a 5-mL HiTrap SP cation-exchange column (GE Healthcare) and eluted with a salt gradient (10–1000 mM NaCl). Peak fractions were concentrated to ~0.5 mL with a 50 kDa concentrator (Amicon Ultra, Millipore), and loaded onto an S200 gel-filtration column (GE Healthcare) in a buffer containing 25 mM HEPES, pH 7.0, 200 mM NaCl, 1 mM dithiothreitol (DTT). Peak fractions were pooled and concentrated to ~15 mg/ml as measured by UV_280_ using a Nanodrop and flash-frozen for storage at −80 °C until use. All protein mutants were generated using the QuikChange protocol from Stratagene and obtained following the expression and purification protocols described above. The primers used to generate the mutants are listed in **Supplementary Table 1**.

### Binary and Ternary *Sc*NatC Expression and Purification

*Sc*Naa30^1-161^ and *Sc*Naa35^19-733^ were cloned into a pET DUET vector containing an TEV cleavable N-terminal His_6_tag. Binary complex (*Sc*Naa30^1-161^/*Sc*Naa35^19-733^) was obtained by transforming this plasmid in ScarabXpress T7lac (Scarab Genomics) competent *E. coli* cells, which were grown in terrific broth media (DOT Scientific) supplemented with ampicillin (100 μg mL^-1^) at 37°C to an OD_600_ of ~0.9 and induced by addition of 0.5 mM IPTG at 17°C for 16 hr. All subsequent purification steps were carried out at 4°C. Cells were isolated by centrifugation, lysed by sonication in lysis buffer containing 25 mM Tris, pH 8.0, 300 M NaCl, 10 mM β-ME and 10 mg/mL PMSF. The lysate was clarified by centrifugation and passed over nickel resin, which was subsequently washed with >20 CV of lysis buffer supplemented with 25 mM imidazole. The protein was eluted in lysis buffer supplemented with 200 mM imidazole by batch elution. TEV protease (~1 mg/ml) was added to the eluted protein and dialyzed into buffer containing 25 mM Tris, pH 8.0, 300 mM NaCl, 5 mM Imidazole, 10 mM β-ME. This solution was passed through an additional nickel column to remove TEV protease as well as any uncut binary *Sc*NatC. The resin was then washed with approximately 3 CV of dialysis buffer supplemented with 25 mM imidazole, which was then pooled with the initial flowthrough. This solution was dialyzed into ion exchange buffer containing 25 mM HEPES, pH 7.5, 50 mM NaCl and 10 mM β-ME and loaded onto a 5 ml HiTrap SP anion exchange column (GE Healthcare). The binary complex *Sc*NatC was then eluted using a salt gradient (50–750 mM NaCl). Peak fractions were concentrated using a 50-kDa MWCO concentrator (Amicon) and further purified Superdex 200 Increase 10/300 GL gel filtration column (GE Healthcare) in a buffer containing 25 mM HEPES, pH 7.0, 200 mM NaCl, 1 mM TCEP. Peak fractions were concentrated to a UV_280_ of ~4-5 mg mL^-1^ as measured by nanodrop. The protein was then flash-frozen and stored at −80°C until use.

To obtain the ternary *Sc*NatC complex, *Sc*Naa38^1-70^ was cloned into a pRSF vector with a TEV-cleavable N-terminal STREP-tag. This plasmid was cotransformed with the pET DUET plasmid containing *Sc*Naa30^1-161^ and *Sc*Naa35^19-733^ and cultured as described above except with the addition of kanamycin (50 μg mL^-1^) to select for pRSF *Sc*Naa38^1-70^ plasmid. The purification of the ternary *Sc*NatC complex was the same as described for the binary *Sc*NatC complex.

### Acetyltransferase Activity Assays

All acetyltransferase assays were carried out at room temperature in a reaction buffer containing 75 mM HEPES, pH 7.0, 120 mM NaCl, 1 mM DTT as described previously (Deng et al., 2019). The full sequence of peptide substrates are listed below: “SASE” peptide (NatA-type): NH_2_-SASEAGVRWGRPVGRRRRP-COOH; “MDVF” peptide (NatB-type): NH_2_-MDVFMKGRWGRPVGRRRRP-COOH; “MLRF” peptide (NatC-type): NH_2_-MLRFVTKRWGRPVGRRRRP-COOH; “MLGP” peptide (NatE-type): NH_2_-MLGPEGGRWGRPVGRRRRP-COOH; “SGRG”/H4 peptide (NatD-type): NH_2_-SGRGKGGKG LGKGGAKRHR-COOH). All peptides were purchased from GenScript. To evaluate *Sp*NatC activity against these peptides, 50 nM *Sp*NatC was mixed with 50 μM Ac-CoA and 500 μM peptide, and allowed to react for 10 min. To determine steady-state catalytic parameters of *Sp*NatC (WT or mutants) with respect to the peptide substrate, 50 nM *Sp*NatC (WT or mutants) was mixed with 50 μM Ac-CoA (^14^C-labeled, 4 mCi mmol^-1^; PerkinElmer Life Sciences) and varying peptide concentrations (ranging from 1.95 μM to 500 μM, 9-data points) for 5-minute reactions. P81 paper was purchased from SVI (St. Vincent’s Institute Medical Research). All radioactive count values were converted to molar units with a standard curve created with known concentrations of radioactive Ac-CoA added to scintillation fluid. GraphPad Prism (version 5.01) was used for all data fitting to the Michaelis–Menten equation. The errors in **Table 2** represent the standard deviation of the best fit values of the curves.

For the comparison of ternary and binary *Sc*NatC complex activity, 10 nM of either binary *Sc*NatC or ternary *Sc*NatC was mixed with 100 μM of ^14^C-labeled Ac-CoA and “MLRF” peptide, for a 12-minute reaction in buffer containing 50 mM HEPES, pH 7.5, 150 mM NaCl and 1 mM DTT.

For the activity comparison of ternary *Sp*NatC and other NATs in the presence or absence of IP_6_, 50 nM of *Sp*NatC, *Sp*NatA or hNatB was mixed with 50 μM of ^14^C-labeled Ac-CoA, 50 μM of their peptide substrate, in reaction buffer containing 75 mM HEPES, pH 7.0, 120 mM NaCl, 1 mM DTT, supplemented with either 0 or 2 μM IP_6_ (Sigma-Aldrich), for a 5-minute reaction. The peptide substates for *Sp*NatA and hNatB are SASE peptide and MDVF peptide, respectively. *Sp*NatA (Liszczak et al., 2013) and hNatB (Deng et al., 2020b) are purified as previously described.

### ITC measurements

ITC measurements were carried out using a MicroCal iTC200 at 20 °C. Samples were dialyzed into buffer containing 25 mM HEPES pH 7.0, 200 mM NaCl, and 1 mM DTT. 15 μM of *Sp*NatC in the cell and 300 μM of IP_6_ in the syringe were used. The raw data were analyzed with the MicroCal ITC analysis software.

### Differential Scanning Fluorimetry Assays

Sypro Orange (50,000X stock, ThermoFisher Scientific) was diluted 1:200, and 4 μL was mixed with 16 μL solution with 0.1 mg/ml of various NatC samples in pH 7.0, 200 mM NaCl, and 1 mM DTT, with or without 10 μM IP_6_. Fluorescent readings were recorded using a qPCR (ABI 7900 RealTime PCR) with a 1% ramp rate, while heated from 20 °C to 95 °C. Melting curves were generated from these readings and melting temperatures were determined by taking the first derivative of the curves. DSF scans of all samples were performed in triplicate as technical replicates. Error bars in the figure indicates the Standard Deviation (SD) of each sample.

### Cryo-EM data Collection

For initial sample screening, 0.05 mg/ml fresh *Sp*NatC sample was prepared with 3-molar excess of both bisubstrate and IP_6_. *Sp*NatC particles on cryo grids exhibited a severe preferred orientation, which generated an incorrect initial 3D model (data not shown). Further screening by addition of detergent NP-40 in sample did not improve this situation. To solve this issue, 1 μL of 80 mM CHAPSO was mixed with 20 μL of 12 mg/mL *Sp*NatC. 3 μL of this mixed sample was applied to Quantifoil R1.2/1.3 holey carbon support grids, blotted and plunged into liquid ethane, using an FEI Vitrobot Mark IV. An FEI TF20 was used for screening the grids and data collection was performed with a Titan Krios equipped with a K3 Summit direct detector (Gatan), at a magnification of 105,000 ×, with defocus values from −0.1 to −3.0 μm. Each stack was exposed and counted in super-resolution mode with a total dose of 42 e^-^/Å^2^, resulting in 35 frames per stack. Image stacks were automatically collected with EPU. A full description of the Cryo-EM data collection parameters can be found in **Table 1**.

### Cryo-EM Data Processing

Original image stacks were summed and corrected for drift and beam-induced motion at the micrograph level using MotionCor2 (Zheng et al., 2017), and binned twofold, resulting in a pixel size of 0.84 A° /pixel. After motion correction in Relion 3.0 (Kimanius et al., 2016), corrected micrographs were imported into CryoSPARC (Punjani et al., 2017) to perform defocus estimation, the resolution range of each micrograph with Gctf (Zhang, 2016) and all the subsequent data analysis. 2D classifications were performed on the particles auto-picked by “blob picker” with particle diameter of 100 −200 Å. The representative 2D classes were used as templates to further auto-pick 1,860,276 particles from 5514 micrographs. After bad particles were removed by two runs of 2D classification, 849,844 particles were used to generate four Ab-Initio models and two rounds of heterogeneous refinement were performed using the four models. 508, 298 particles were used for auto refinement and per particle CTF refinement. The final map was refined to an overall resolution of 3.20 Å, with local resolution estimated in Cryo-SPARC (Punjani et al., 2017).

### Cryo-EM Model Building and Refinement

The *Sp*NatC atomic model was manually built de novo using the program COOT (Emsley and Cowtan, 2004) according to the Cryo-EM map, with the guidance of predicted secondary structure and bulky residues such as Phe, Tyr, Trp and Arg. The complete model was then refined by real-space refinement in PHENIX (Adams et al., 2010). All representations of Cryo-EM density and structural models were performed with Chimera (Pettersen et al., 2004) and PyMol (Schrodinger, 2015) (https://pymol.org/2/). The sequence alignments with secondary structure display were created by ESPript 3.0 (Robert and Gouet, 2014). The surface area calculation was performed using PDBePISA (Krissinel and Henrick, 2007) (Proteins, Interfaces, Structures and Assemblies) (http://www.ebi.ac.uk/pdbe/pisa/).

## ACKNOWLEDGEMENTS

This work was supported by NIH grant R35 GM118090 awarded to R.M and R01 NS103873 awarded to E.J.P. We thank Zuo Biao and Sudheer Molugu from the University of Pennsylvania Electron Microscopy Resource Lab for help with initial cryo-grids screening; and Stefan Steimle from the Beckman Center for Cryo-EM at the University of Pennsylvania for technical assistance on data collection. We also thank Elaine Zhou for help with analyzing the DSF data.

## Author Contribution

Conceptualization, S.D., B.P., J.S., L.G., X. W., E.J.P., and R.M.; Methodology, S.D., B.P., J.S., L.G., X. W., E.J.P., and R.M..; Investigation, S.D., B.P., J.S., L.G., X. W.; Formal Analysis, S.D., L.G., and R.M.; Writing – Original Draft, S.D.; Visualization, S.D.; Writing – Review and Editing, S.D., L.G., B.P., J.S., X. W., E.J.P., and R.M.; Funding Acquisition, E.J.P., R.M.; Resources, R.M.; Supervision, E.J.P. and R.M.

## Data availability

The cryo-electron microscopy map for the *Sp*NatC – Bisubstrate analogue-IP_6_ and the atomic coordinate have been deposited in the EMDataBank and ProteinDataBank, with accession codes EMD-23110 and PDB: 7L1K, respectively. Primary data are available upon reasonable request from the corresponding author.

## Supplementary information

**Supplementary Table 1.**
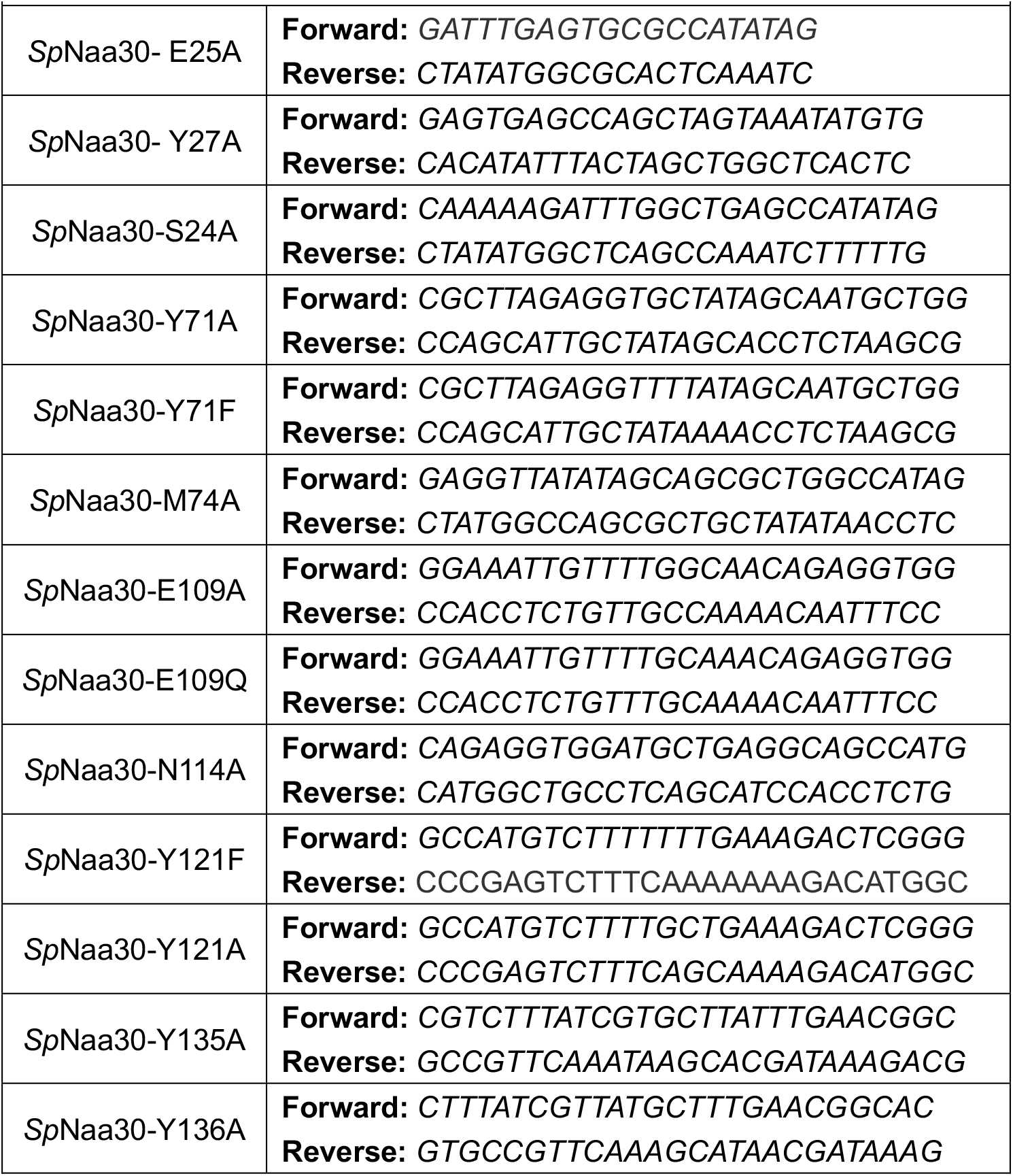
Sequence of primers for preparing mutations.

**Supplementary figure 1.**
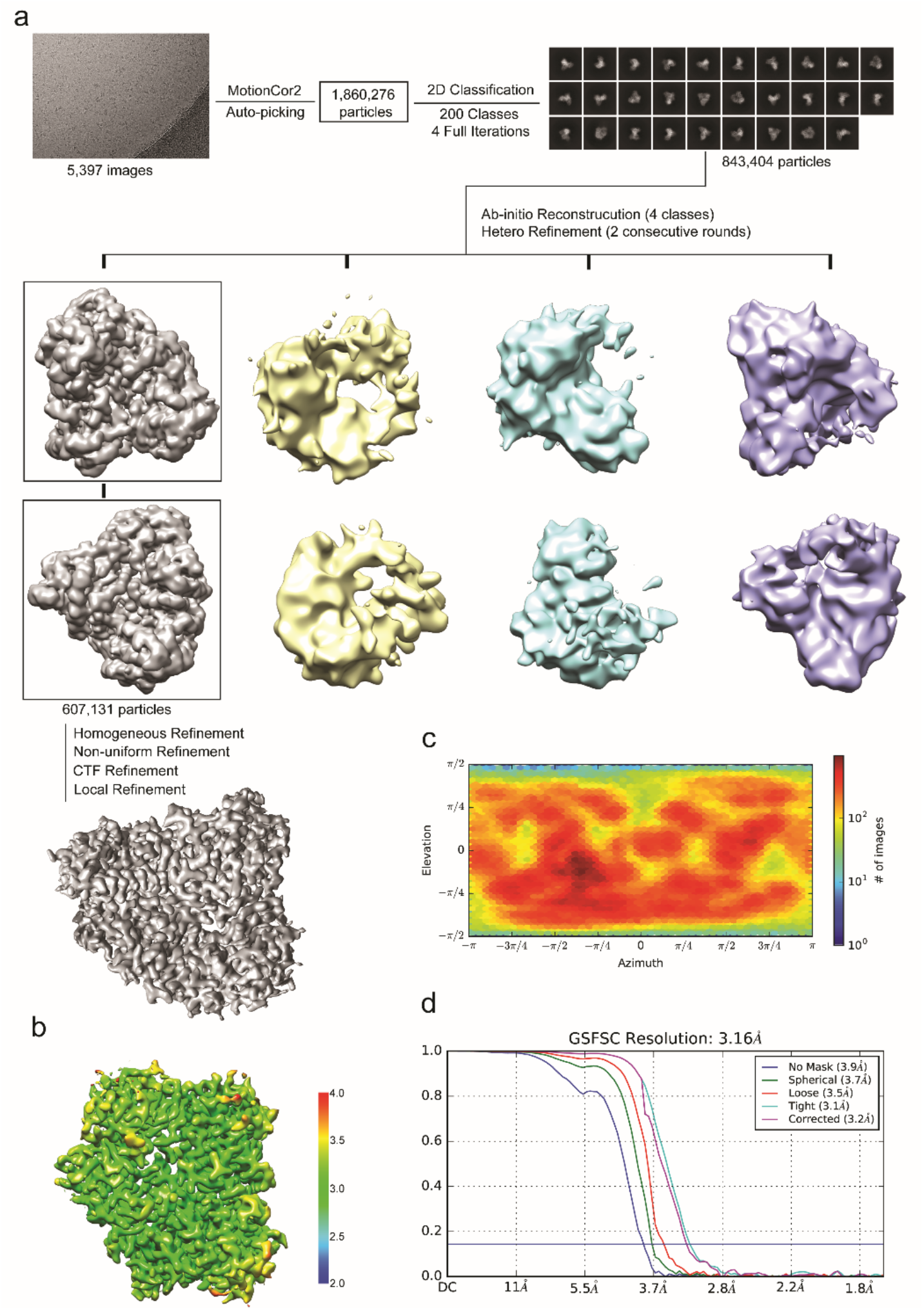
Cryo-EM workflow and resolution of *Sp*NatC EM map. (**a**) 2D and 3D classification scheme and workflow for *Sp*NatC EM map determination. (**b**) Local resolution of the final *Sp*NatC EM map. (**c**) Viewing direction distribution of *Sp*NatC final EM map 3D reconstruction generated by cryoSPARC v2. (**d**) Gold standard Fourier Shell Correlation (FSC) curves of *Sp*NatC EM map 3D reconstruction, generated by cryoSPARC v2.

**Supplementary figure 2.**
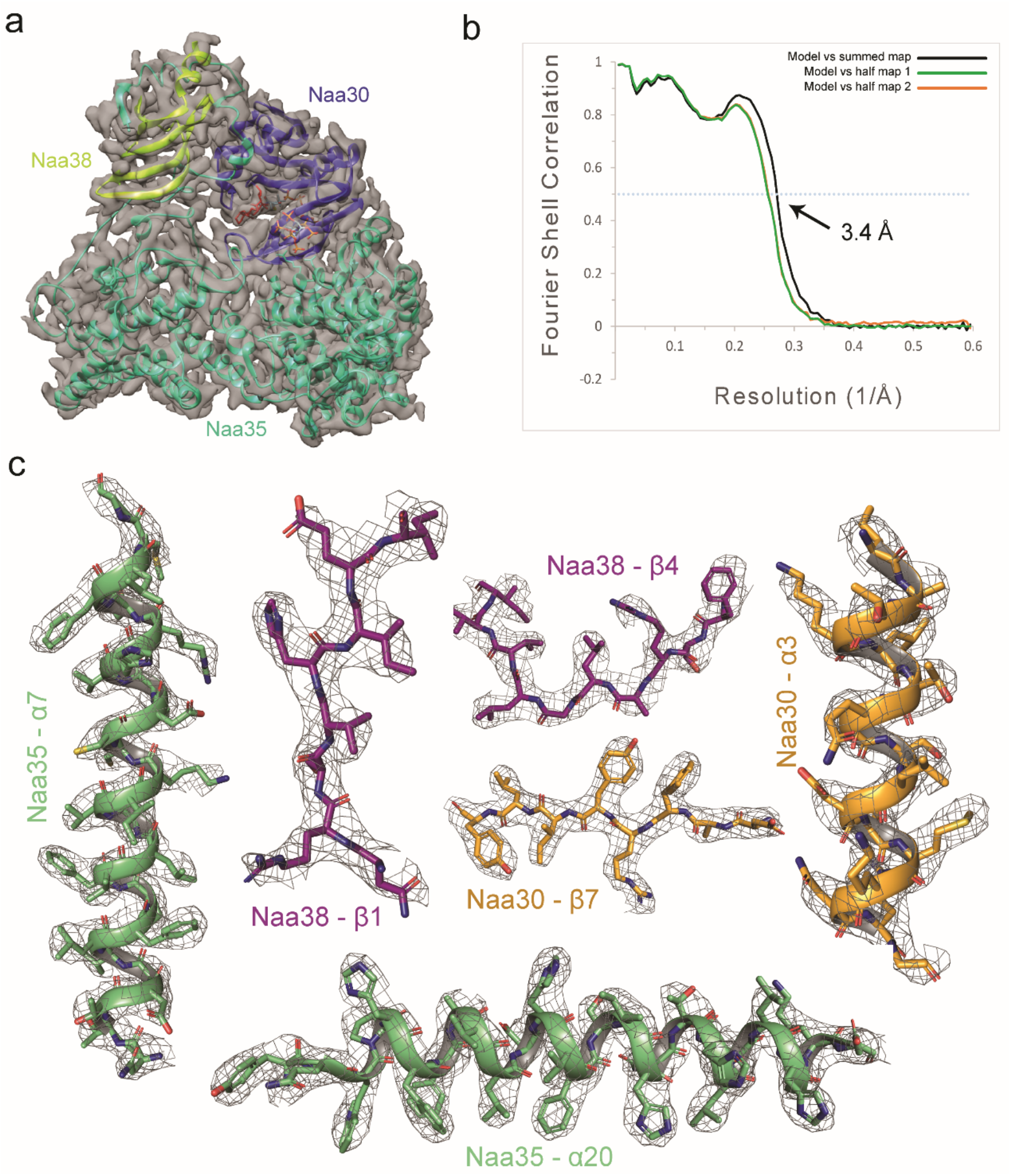
Correlation between Cryo-EM map of *Sp*NatC EM and its atomic model. **(a)** Atomic model of *Sp*NatC fitted into the Cryo-EM map. **(b)** FSC curves of the refined model versus the overall map that it was refined against (black); of the model refined in the first of the two independent maps used for the gold-standard FSC versus the same map (green); and of the model refined in the first of the two independent maps versus the second independent map (orange). **(c)** The fit of several helical segments or β-strands from all three subunits of NatC in the EM density. The contour level is 5 sigma.

**Supplementary figure 3.**
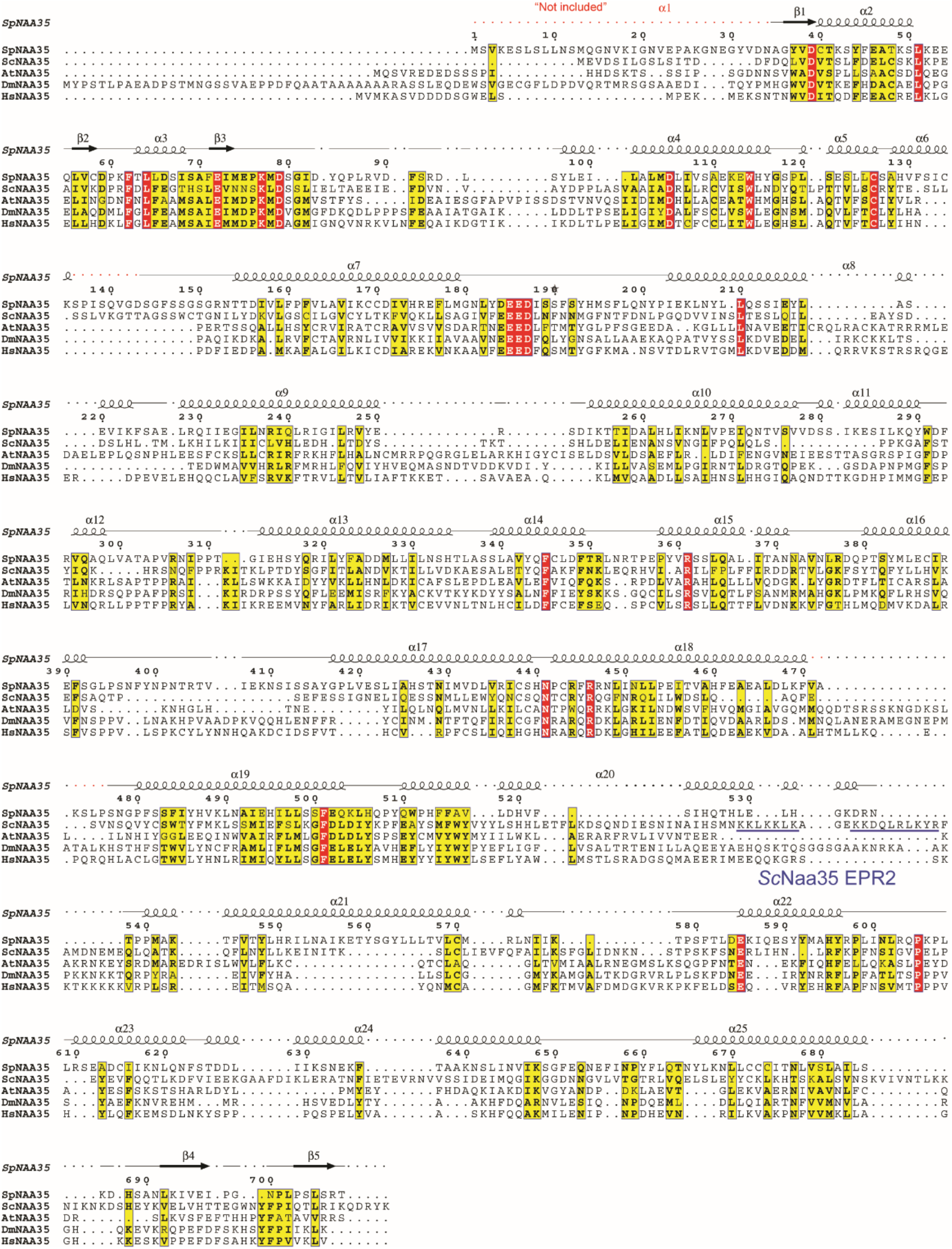
Sequence alignment of Naa35 orthologs from *Schizosaccharomyces pombe* (Sp), *Saccharomyces cerevisiae (*Sc), *Arabidopsis thaliana* (At), *Drosophila melanogaster (*Dm), and *Homo sapiens* (Hs). Numbering and secondary structure elements for *Sp*Naa30 are indicated above the sequence alignment. Residues truncated from the *Sp*Naa35 protein construct used in this study are indicated above in red (“Not included”). EPR2 (**Figure 5a)** from *Sc*Naa35 is annotated in blue.

**Supplementary figure 4.**
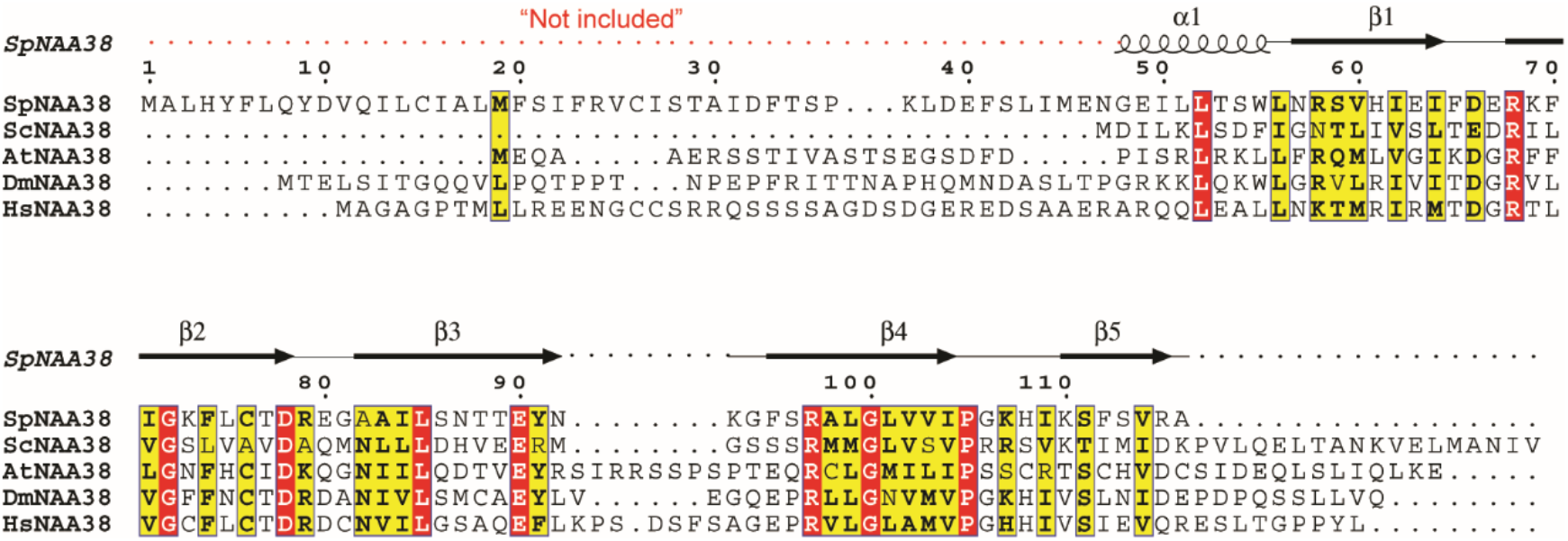
Sequence alignment of Naa38 orthologs from *Schizosaccharomyces pombe* (Sp), *Saccharomyces cerevisiae* (Sc), *Arabidopsis thaliana* (At), *Drosophila melanogaster (*Dm), and *Homo sapiens* (Hs). Numbering and secondary structure elements for *Sp*Naa38 are indicated above the sequence alignment. Residues truncated from the *Sp*Naa38 protein construct used in this study are indicated above in red (“Not included”).

**Supplementary figure 5.**
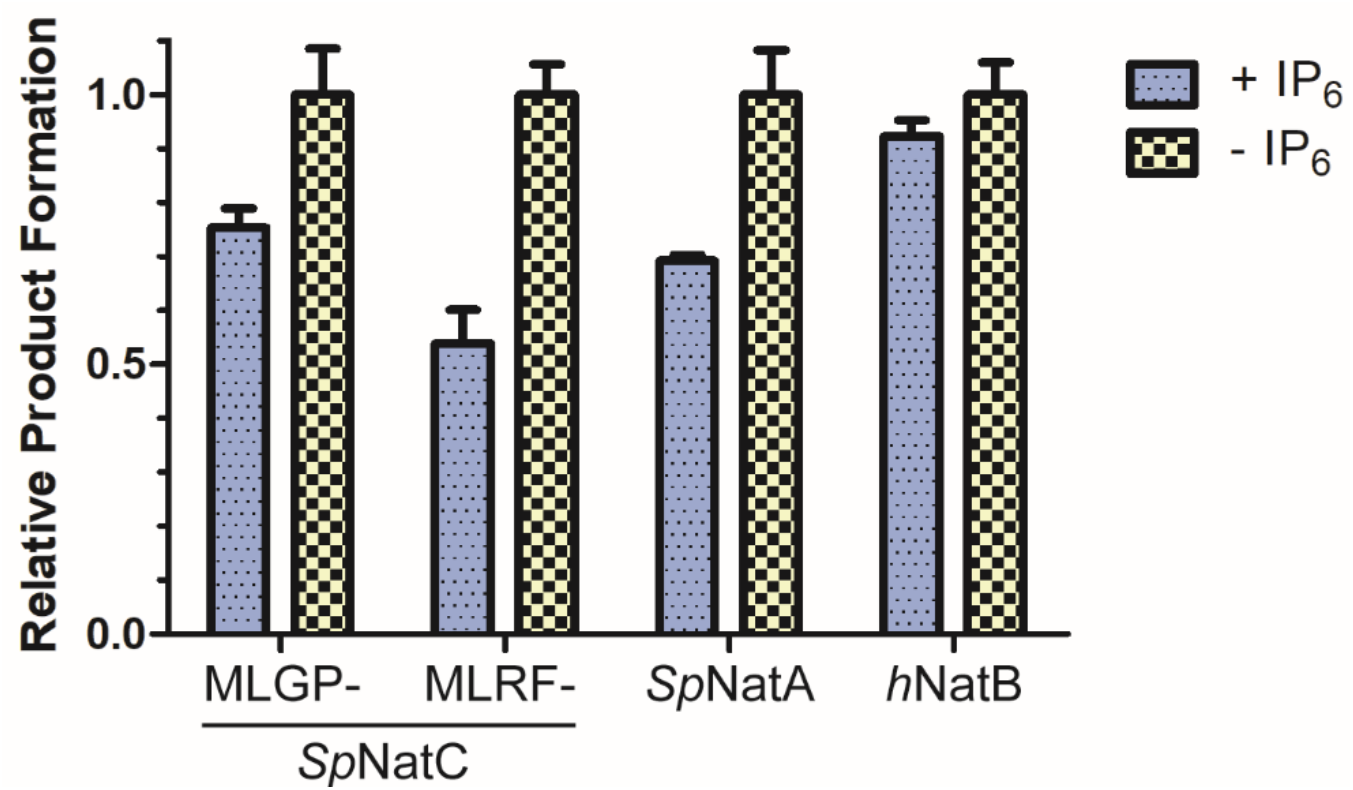
The activity impacts of IP_6_ on selective NATs. 50 nM of either *Sp*NatC, *Sp*NatA, and hNatB was mixed with 2 μM or no IP_6_ for measuring production formation. Data was normalized to NATs activity without IP_6_. Error bars indicates Mean with SEM, n=3.

